# Comprehensive ctDNA Profiling Enables Tissue-of-Origin Prediction and Actionable Biomarker Detection in Cancer of Unknown Primary

**DOI:** 10.64898/2026.06.18.732338

**Authors:** Jana Löptien, Maximilian Haas, Maria Pouyiourou, Christian Müller, Cornelia Coith, Tilmann Bochtler, Mingyang Cai, Elmira Forouzmand, Yupeng He, Olaf Neumann, Albrecht Stenzinger, Sabine Riethdorf, Alwin Krämer, Klaus Pantel, Harriet Wikman

## Abstract

Most patients with cancer of unknown primary (CUP) still receive platinum-based chemotherapy and have a poor prognosis, with overall survival of less than one year. Recent studies suggest improved outcomes with molecularly guided or site-specific therapies informed by molecular tissue profiling. Here, we analyzed ctDNA from 190 CUP patients using an integrated genomic and epigenomic assay to identify actionable alterations and predict tissue-of-origin (ToO). Integration of actionable biomarkers, ToO prediction and clinical data yielded diagnostic, prognostic or therapeutic information in 90% of unfavorable CUP cases and 88% of patients analyzed at first diagnosis. High ctDNA tumor fraction was associated with poorer prognosis in both favorable and unfavorable CUP. These findings highlight the clinical utility of ctDNA analysis for therapeutic decision-making in CUP and support its incorporation into the diagnostic work-up, particularly when tissue samples are unavailable or insufficient for molecular testing.

## Introduction

Cancer of unknown primary (CUP) is a metastatic malignancy in which the site of the primary tumor remains unidentified despite extensive diagnostic evaluation. It represents approximately 1-3% of all cancer diagnoses, and is associated with a poor prognosis with a 1-year survival rate of less than 20% [1–3]. CUP is categorized into favorable and unfavorable subtypes, with 80-85% of patients classified into the unfavorable subtype [4].

The favorable subtype includes cases with single-site or oligometastatic disease that are amenable to local ablative treatment, and tumors that resemble a defined cancer entity and can be treated with site-specific therapies [4]. Patients within this subgroup generally have a better prognosis. In contrast, patients with unfavorable CUP, who do not meet the criteria for one of the favorable categories, have a median overall survival (OS) ranging from 6 to 10 months only [3,4]. For decades, treatment for unfavorable CUP patients relied on platinum-based combination chemotherapy, with limited success. This underscores the aggressive nature of CUP and highlights the need for comprehensive characterization of its mutational and epigenetic profiles as basis for the identification of targeted therapeutic approaches and tissue-of-origin (ToO) prediction strategies, respectively, to guide treatment decisions as alternatives to non-specific chemotherapy [5–8].

The molecular landscape of CUP reflects its marked heterogeneity, with each patient exhibiting a distinct mutational profile. Approximately one-third to one-half of CUP cases harbor at least one potentially actionable genomic alteration [9–12]. The CUPISCO trial demonstrated that unfavorable CUP patients benefit from targeted therapies aimed at actionable alterations, further supporting comprehensive genomic profiling in CUP. Overall, current research into precision oncology approaches in CUP is evolving rapidly. Current studies suggest that both molecularly-guided tumor-agnostic and site-specific treatment strategies may improve clinical outcomes compared to empirical chemotherapy, supporting the continued development of individualized treatment approaches in this challenging disease [5,7,13].

Most studies to date have characterized the genomic landscape of CUP using tissue biopsies [3]. However, in clinical practice, tumor tissue is often limited and already exhausted when it comes to mutational- or methylation-based analysis. This is due to the extensive diagnostic immunohistology procedures required to identify or exclude a putative primary site as well as the usually small biopsies taken in the absence of a primary tumor that can be operated on [4]. In this context, liquid biopsy approaches, particularly comprehensive genomic profiling (CGP) of circulating tumor DNA (ctDNA), offer a valuable complementary strategy for diagnosis and clinical decision-making [10,14,15]. Compared to tissue biopsies, liquid biopsies are minimally invasive, more easily accessible, and capable of capturing the molecular heterogeneity of the disease across multiple metastatic sites simultaneously. Additionally, in contrast to tissue biopsies, they are well suited for frequent longitudinal monitoring and routine follow-up [16,17].

In this study, we used a blood-based comprehensive genomic and epigenomic sequencing approach to investigate the plasma genomic landscape of 190 CUP patients and analyze the utility of ctDNA to detect actionable alterations, determine the ToO, and assess the prognostic impact of ctDNA analysis.

## Results

### Tumor-agnostic liquid biopsy-based molecular profiling of CUP

Of the 190 plasma samples analyzed using the Guardant360 Liquid assay (Guardant Health, Palo Alto, CA, USA), 187 (98%) met quality-control criteria and were included in subsequent analyses (Figure 1a). In each sample, the circulating free DNA (cfDNA) yield was above the minimally recommended input of 5 ng (median 25.5 ng, range 5.7-700.1 ng; Figure S1a). Somatic non-synonymous mutations, including single nucleotide variants (SNVs), small insertions or deletions of nucleotides (indels), copy number variations (CNVs), and fusions were detected in 98% (183/187) of patients. The median number of mutations per patient was 9 (range: 0-371; Figure 1b). The number of mutations per patient correlated with cfDNA input (Spearman’s rank correlation coefficient, p<0.0001, rho=0.4033; Figure S1b).

**Figure 1:**
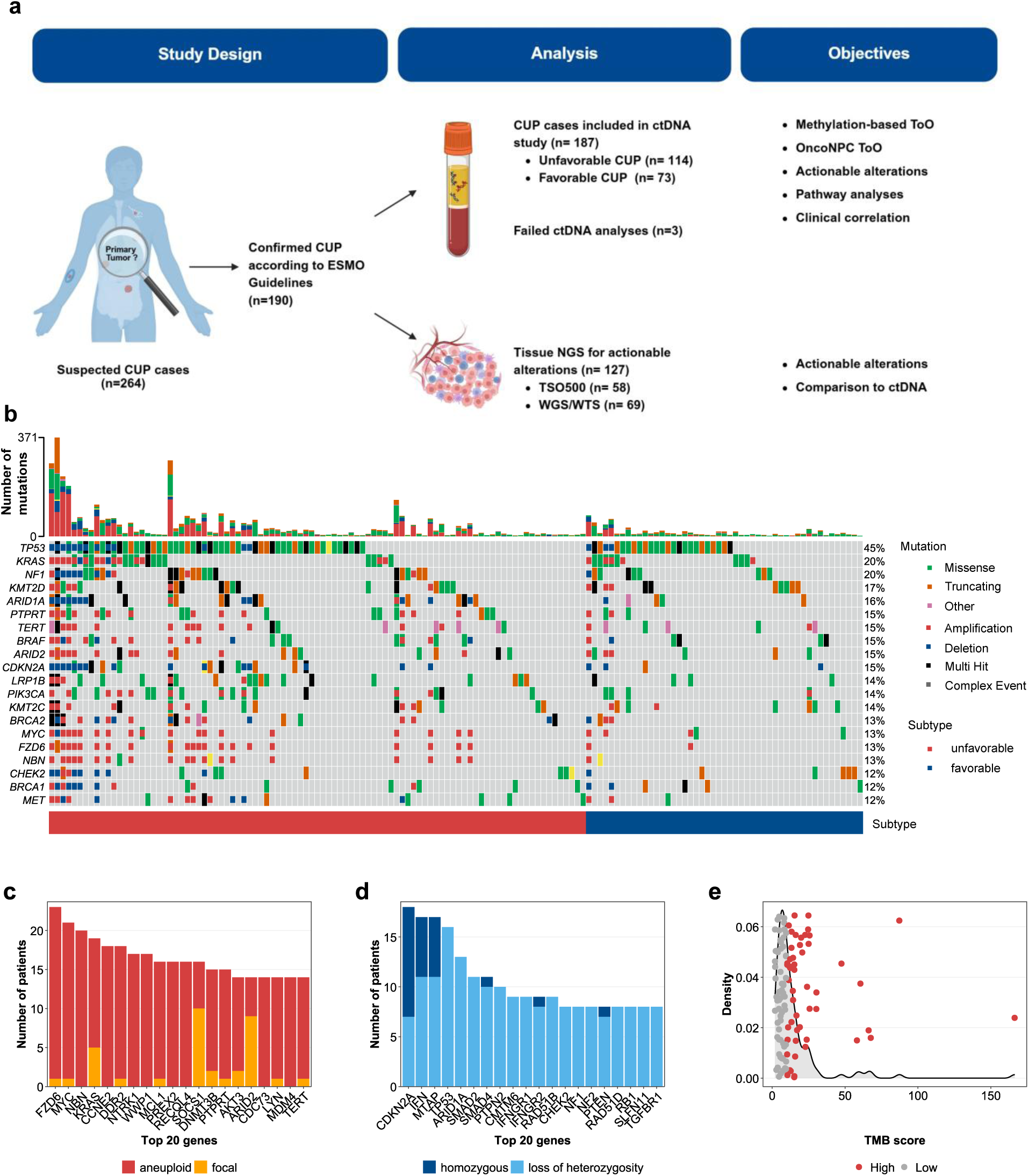
Deep sequencing enables mutation and methylation profiling of ctDNA for the majority of CUP patients. **(a)** Flow chart showing the study workflow and objectives. **(b)** Oncoplot showing the top 20 mutated genes in all samples, including a bar plot representing the number of mutations per patient. **(c,d)** Bar plots showing the top 20 genes harboring focal (high-level) and aneuploid (low-level) amplifications or homozygous deletions and loss of heterozygosity, respectively. **(e)** Density plot showing the distribution of patients by TMB score using Kernel density estimates (curve) overlaid with individual patient TMB scores (x-axis) distributed randomly along the y-axis (points). Samples with high TMB (>10 mut/Mb) are shown in red.

In line with previous molecular profiling studies of CUP liquid and tissue biopsies, the five most frequently altered genes were *TP53* (45%), *KRAS* (20%), *NF1* (20%), *KMT2D* (17%), and *ARID1A* (16%; Figure 1b) [12,14,18]. CNVs were detected in 71 patients. Of the top 20 genes with amplifications, *KRAS* and *MYC* were commonly described in CUP tissue before [12,14,18]. In our study focal high-level amplifications were most commonly detected in *SOCS1* (5%) and *ARID2* (5%; Figure 1c). The most common homozygous deletion was found at 9p21.3, eliminating the *CDKN2A* gene in 6% (11/187) of patients with co-occurring deletion of the type I interferon gene cluster (IFN) and MTAP in six patients (Figure 1d), as described before [19].

High tumor mutational burden (TMB, defined as ≥ 10 mut/Mb) was detected in 51 patients (27%), while the overall median of TMB was 9.3 mut/Mb (range 0-166.41; Figure 1e). TMB did not correlate with the cfDNA input (Spearman’s rank correlation coefficient: p=0.6355, rho=0.0447; Figure S1d). Microsatellite-instability (MSI) was detected in five patients (Figure S1e), and homologous recombination deficiency (HRD) was found in only two patients (Figure S1f).

A methylation-based tumor fraction (EpiTF) was calculated, which has been shown to closely correlate with the tumor burden [20]. Here, the EpiTF showed a median of 0.32% tumor-derived DNA in the plasma (range: 0.00-83.36%, Figure S1g). Similar to the number of mutations, the EpiTF correlated significantly, with the cfDNA input (Spearman’s rank correlation coefficient: p<0.0001, rho= 0.3964; Figure S1c), as well as with the number of mutations (Spearman’s rank correlation coefficient: p<0.0001, rho=0.796; Figure 2a), indicating a higher detection rate in high shedding tumors, as described before [21].

**Figure 2:**
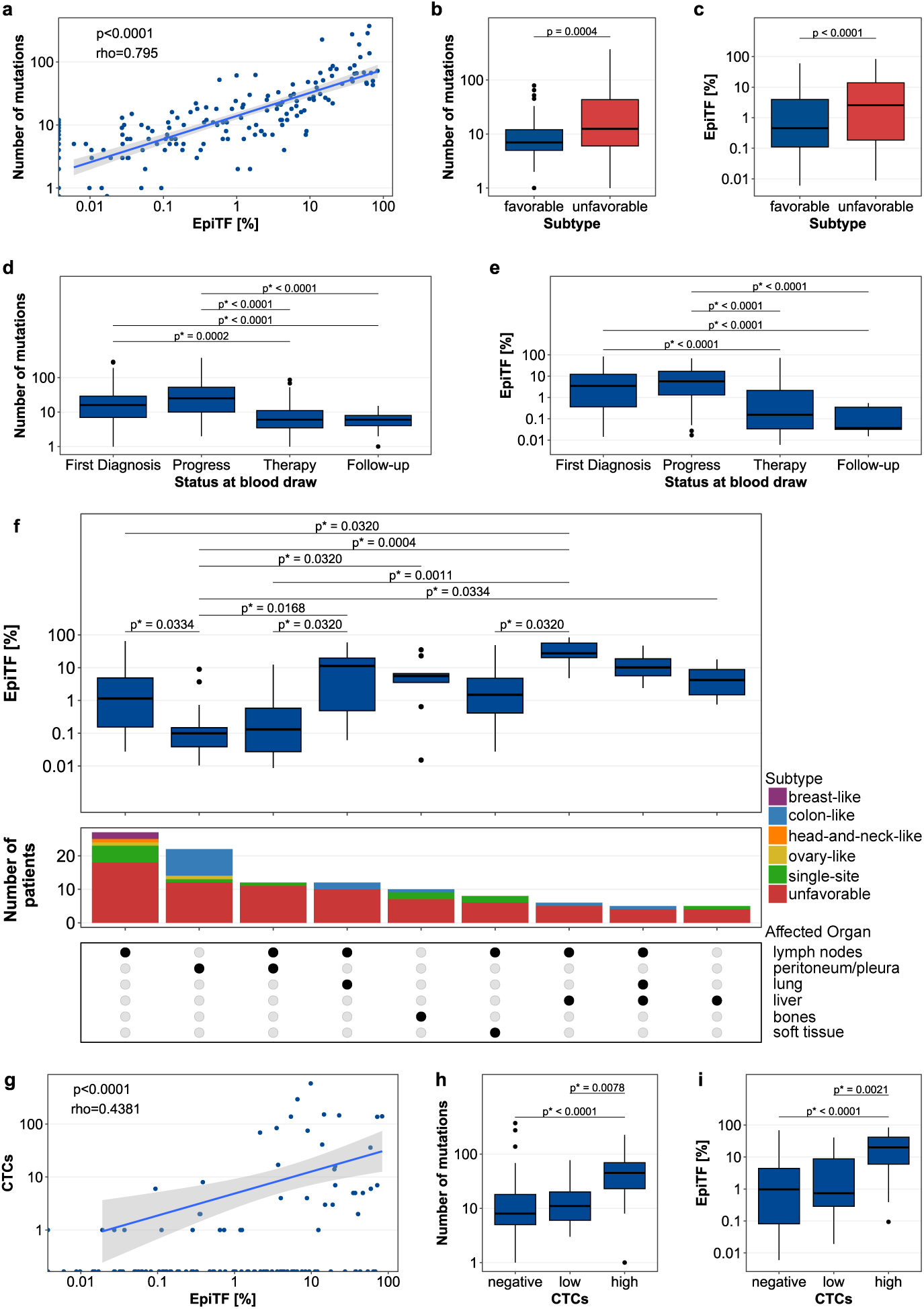
Blood-based tumor fraction and number of mutations correlate with clinical features. **(a)** Correlation of the number of mutations with the epigenetic tumor fraction (EpiTF), Spearman’s rank correlation: p<0.0001, rho=0.795. The points represent individual patients; the line depicts the linear regression fit, and the shaded grey area represents the 95% confidence interval. **(b,c)** Boxplots showing the association of CUP subtype with the number of mutations or EpiTF, respectively. Wilcoxon rank sum test: p=0.0004 and p<0.0001, respectively. **(d,e)** Boxplots showing the association between blood draw timing and the number of mutations or EpiTF, respectively. Dunn’s test for multiple comparisons with FDR correction. **(f)** Upset-plot depicting the association of affected organs and EpiTF. Bottom: intersection matrix, with filled circles depicting the combination of affected organs in that group (x-axis), middle: bar plot showing the number of patients per intersection group, top: boxplots showing the EpiTF of each intersection group. Dunn’s test for multiple comparisons with FDR correction. **(g)** Correlation of the number of CTCs/7.5 mL blood with EpiTF. Spearman’s rank correlation: p<0.0001, rho=0.4381. The points represent individual patients; the line depicts the linear regression fit, and the shaded grey area represents the 95% confidence interval. **(h,i)** Boxplots showing the association of CTC frequency grouped in negative (CTCs=0), low (CTCs=1-4) and high (CTCs≥5) with the number of mutations or EpiTF, respectively. Dunn’s test for multiple comparisons with FDR correction. For all boxplots, the center line indicates the median, boxes represent the 1^st^ and 3^rd^ quartiles, whiskers extend to 1.5 x the interquartile range (IQR), and points depict outliers of 1.5xIQR.

### Unfavorable subtype is associated with increased rates of molecular alterations

Liquid biopsy analysis revealed an association of both the EpiTF, and the number of mutations detected with several clinical features. In patients with unfavorable CUP, the number of mutations and EpiTF were elevated compared to the favorable subtype (Wilcoxon Rank Sum Test, p=0.0004, p<0.0001, respectively; Figure 2b,c), while there was no difference in TMB observed between the subtypes (Wilcoxon Rank Sum Test, p=0.0677; Figure S2a). As described for ctDNA analysis before [22], we also detected a strong influence of the timepoint of blood collection on the tumor fraction and the number of mutations detected. Patients who received a liquid biopsy assessment before treatment (first diagnosis), or during disease progression, had a significantly higher number of mutations and EpiTF compared to patients who had a liquid biopsy during therapy or during follow-up after therapy completion (Dunn’s test for Multiple Comparisons with FDR correction; p* values see Figure 2d,e). This was not observed for TMB (Kruskal-Wallis Rank Sum test, p = 0.2234; Figure S2b).

Interestingly, the involvement of lungs, bones, and liver was associated with higher EpiTF compared to involvement of peritoneum, pleura, and lymph nodes. (Dunn’s test for Multiple Comparisons with FDR correction; p* values see Figure 2f). In contrast, there was no significant difference in EpiTF, number of mutations and TMB between histologic subtypes (Kruskal-Wallis Rank Sum test, p=0.1845, p=0.4032, p=0.2448, respectively; Figure S2c-e).

Previously, we could show that the detection of circulating tumor cells (CTCs) in whole blood is a strong, independent predictor of survival in patients with CUP [23]. In 44 (28%) out of 158 patients, in whom ctDNA was analyzed in the present study, CTCs were detected. The number of CTCs correlated with the EpiTF and the number of mutations in ctDNA (Spearman’s rank correlation coefficient; p<0.0001 rho=0.4381, p<0.0001, rho=0.3759, respectively; Figure 2g and S2f). Accordingly, the number of mutations and EpiTF were significantly higher in patients with positive CTC status (Wilcoxon Rank Sum test, p<0.0001 for both; Figure S2g,h). This association was even stronger in patients with a high number of CTCs (≥5 CTCs / 7.5 mL; Dunn’s test for Multiple Comparisons with FDR correction, respectively; p* values see Figure 2h,i), which we had shown to be prognostically relevant for OS [23].

### Detection of molecular targets in blood-based mutational profiles

As previous studies have shown improved PFS and OS for patients receiving molecularly-guided therapy based on tissue NGS [5,7], we determined whether ctDNA NGS can detect actionable alterations annotated using OncoKB™ Knowledge Base [24,25]. We detected actionable mutations in 50 patients (27%) and 20 different genes. The most commonly altered actionable genes were *PIK3CA*, *ERBB2*, *MET*, *PTEN*, and *BRAF,* similar to the actionable alterations described in tissue-based assays of CUP patients [12]. Additionally, 51 patients (27%) had at least one actionable biomarker finding, including TMB-High, MSI-High, or HRD [8]. In total, 76 (41%) patients would have therefore been potentially eligible for targeted treatment based on OncoKB™ level of evidence 1 and 2, defined as FDA-approved drugs and standard of care (SOC), respectively (Figure 3a,b) [6]. Importantly, 72% (55/76) of those patients belong to the unfavorable subtype.

**Figure 3:**
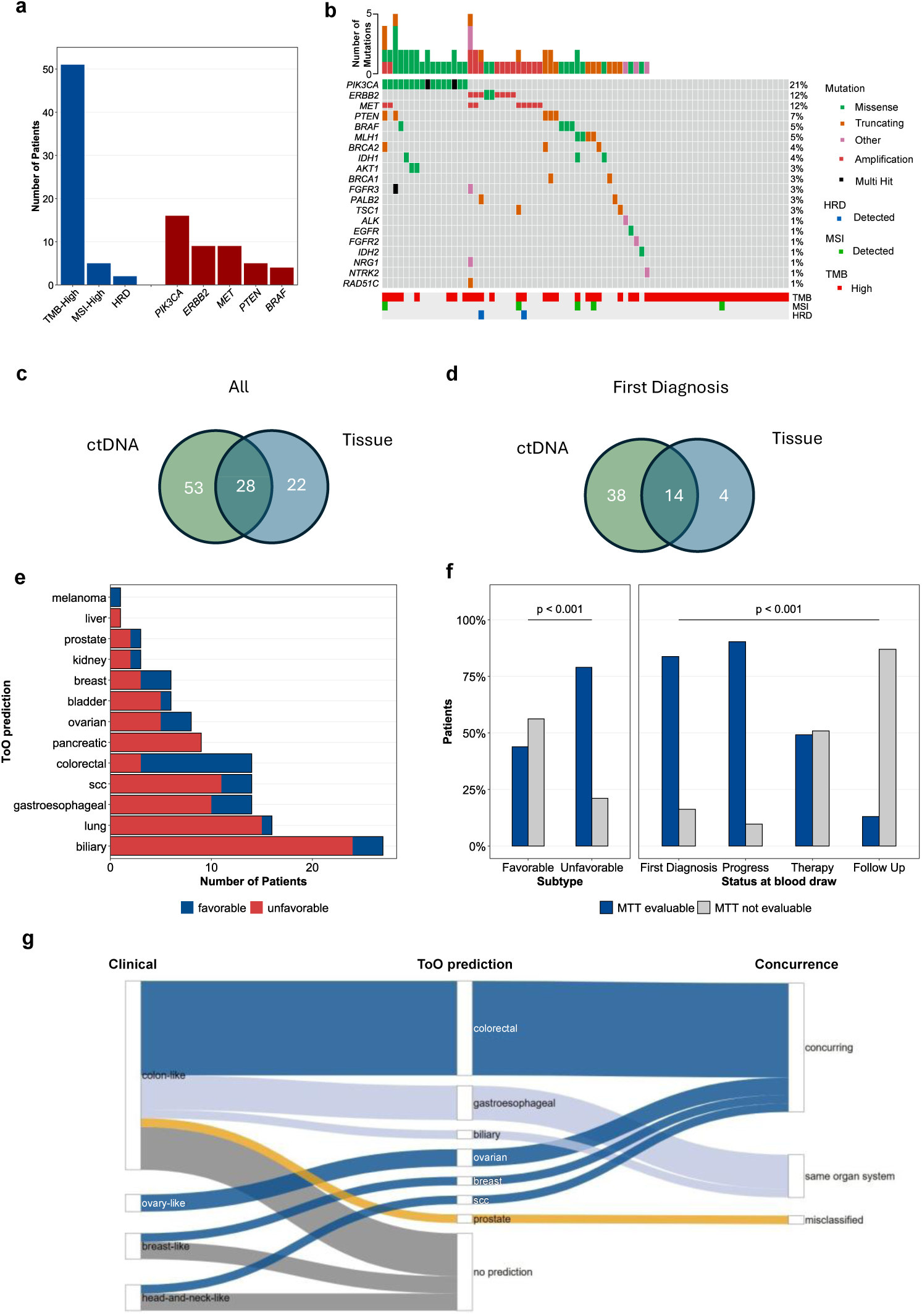
Liquid biopsy enables detection of actionable alterations and ToO prediction. **(a)** Bar plot showing the frequency of actionable alterations of the three biomarkers (TMB-High, MSI-High and HRD) and in the top 5 genes detected. **(b)** Oncoplot showing the actionable alterations detected in individual patients, top: bar plot showing the number of actionable mutations detected per patient. **(c,d)** Venn-diagrams showing the overlap of actionable alterations detected in liquid and tissue biopsy for all patients (n=65) and patients that received both sequencing approaches before treatment (first diagnosis n=29), respectively. **(e)** Column plot showing the proportions of ToO predictions analyzed using MTT. **(f)** Bar plot showing the frequencies of positive ToO results by subtypes (left) and statuses at blood draw (right), Fisher’s exact test, p<0.001 for both. **(g)** Sankey plot showing the ToO predictions for patients belonging to one of the favorable subtypes that resemble a tumor with known primary, and the concordance between the clinically defined potential primary site and the ToO predicted by the MTT algorithm.

To determine the utility of ctDNA analysis for the detection of actionable alterations, we compared ctDNA results to tissue NGS. Of the 127 patients that received tissue NGS, 68 (54%) tumors harbored actionable alterations. Consistent with the ctDNA analysis, the most common genes affected were *PIK3CA* (18/68), *ERBB2* (12/68), and *PTEN* (9/68). To conduct a comparative analysis between tissue and liquid biopsy results, we filtered mutations for genomic locations that were common to both tissue and blood analysis. In total, 103 actionable alterations were found in liquid and/or tissue biopsies of 65 patients. Thereof, 28 (27%) alterations in 26 patients were found in both tissue and liquid biopsy, while 53 (51%) alterations in 25 patients and 22 (21%) alterations in 14 patients were exclusive to liquid and tissue biopsy, respectively. In patients who were untreated before both liquid and tissue biopsy were taken (n=29), 56 actionable alterations were detected. Of these, 68% were exclusively detected in liquid biopsy across 14 patients, 7% exclusively in tissue biopsy across 2 patients, and 25% in both specimen types across 13 patients. These findings suggest that ctDNA sequencing can provide complementary and, in some cases, additional actionable information beyond tissue-based sequencing and can guide clinical decision making for a broader subset of patients. As this effect was most evident at first diagnosis, these findings support performing liquid biopsy early in the diagnostic work-up, at the time of disease onset (number of alterations: Figure 3c,d; number of patients: Figure S3a, b).

### ctDNA-based tissue of origin (ToO) prediction to guide clinical decision making

As identification of the tissue of origin (ToO) and subsequent guided treatment might improve patient outcomes [3,7], we performed methylation-based Molecular Tumor Typing (MTT) of the plasma samples. In our cohort, 13 of 14 different cancer types were predicted in 65% (122/187) of patients. Of those, 75% had a high confidence-level primary prediction. The most common primary predictions were biliary 22% (27/122) and lung 13% (16/122), followed by gastroesophageal, squamous-cell carcinomas (SCC), and colorectal (11%, 14/122 each; Figure 3e).

An evaluable MTT result that generated a ToO prediction was significantly associated with several clinical features. ToO evaluable rate is significantly higher among patients with an unfavorable subtype (79%) and among patients whose blood draw occurred at first diagnosis or disease progression (84% and 90%, respectively; Fisher’s Exact Test, p<0.001 for both; Figure 3f). Furthermore, high ToO evaluable rate was significantly associated with high EpiTF, positive CTC status, TMB-high and cfDNA-high status (Table 1), most likely associated with a higher tumor load.

**Table 1:** Summary Table showing the yield of positive and negative ToO prediction call rate stratified by clinical and molecular features. Fisher’s Exact Test.

| Characteristic | Overall<br>n = 187 | positive ToO call<br>n = 122 | negative ToO call<br>n = 65 | p-value |
| --- | --- | --- | --- | --- |
| **Sex** |  |  |  | 0.2 |
| Female | 108 | 66 (61%) | 42 (39%) |  |
| Male | 79 | 56 (71%) | 23 (29%) |  |
| **Histology** |  |  |  | 0.4 |
| Adenocarcinoma | 127 | 86 (68%) | 41 (32%) |  |
| Squamous-cell carcinoma | 39 | 21 (54%) | 18 (46%) |  |
| Undifferentiated carcinoma | 10 | 7 (70%) | 3 (30%) |  |
| Other | 11 | 8 (73%) | 3 (27%) |  |
| **Status at blood draw** |  |  |  | <0.001 |
| First Diagnosis | 74 | **62 (84%)** | 12 (16%) |  |
| Progress | 31 | **28 (90%)** | 3 (9.7%) |  |
| Therapy | 59 | 29 (49%) | 30 (51%) |  |
| Follow Up | 23 | 3 (13%) | 20 (87%) |  |
| **Subtype** |  |  |  | <0.001 |
| Favorable | 73 | 32 (44%) | 41 (56%) |  |
| Unfavorable | 114 | **90 (79%)** | 24 (21%) |  |
| **EpiTF** |  |  |  | <0.001 |
| negative | 46 | 0 (0%) | 46 (100%) |  |
| Low (< 1.5%) | 71 | 53 (75%) | 18 (25%) |  |
| High (≥ 1.5%) | 70 | **69 (99%)** | 1 (1.4%) |  |
| **CTC status** |  |  |  | <0.001 |
| Negative | 113 | 66 (58%) | 47 (42%) |  |
| Positive | 42 | **39 (93%)** | 3 (7.1%) |  |
| **TMB** |  |  |  | <0.001 |
| Low (< 10 mut/Mb) | 66 | 45 (68%) | 21 (32%) |  |
| High (≥ 10 mut/Mb) | 51 | **49 (96%)** | 2 (3.9%) |  |
| **cfDNA** |  |  |  | 0.001 |
| low (5-17.74 ng) | 62 | 31 (50%) | 31 (50%) |  |
| medium (17.74-30 ng) | 48 | 30 (63%) | 18 (38%) |  |
| high (> 30 ng) | 77 | **61 (79%)** | 16 (21%) |  |

For patients belonging to one of the favorable CUP subtypes that resemble a defined cancer entity, 70% (21/30) received a primary prediction. In only one patient the ToO prediction differed completely from the respective favorable CUP subtype, with a clinical colon-like CUP predicted as prostate cancer. The other samples had either concordant predictions or predictions of related organ systems (Figure 3g).

To determine if mutational analysis further increased the ToO prediction frequency, we used OncoNPC [26] as a mutation-based ToO prediction algorithm and reported predictions with a confidence score >0.8. Mutation-based ToO predictions were obtained in 16% of patients (30/187), which was significantly lower than that of methylation-based ToO prediction (Fisher’s exact test, p<0.0001). The majority (19/30, 63%) of mutation-based ToO predictions were either concordant or of additional value compared with the methylation-based ToO prediction (Supplementary Table S1). For two patients where methylation-based ToO was not evaluable, a mutation-based ToO was administered. Discrepancies between mutation-based and methylation-based ToO predictions were observed in nine patients and can likely be attributed to the limited robustness of mutational signatures using panel sequencing.

To assess the concordance between methylation-based ToO predictions and known primary tumors, we performed unsupervised clustering by generating a UMAP using methylation data from liquid biopsies. For this, our CUP cohort was compared to 34,911 advanced stage cancer patients with a known primary (Figure 4a). The UMAP showed that most CUP cases indeed clustered together with samples from primary cancer types matching the ToO prediction. This indicates that the methylation-based ToO predictions can accurately predict the putative primary tumor type.

**Figure 4:**
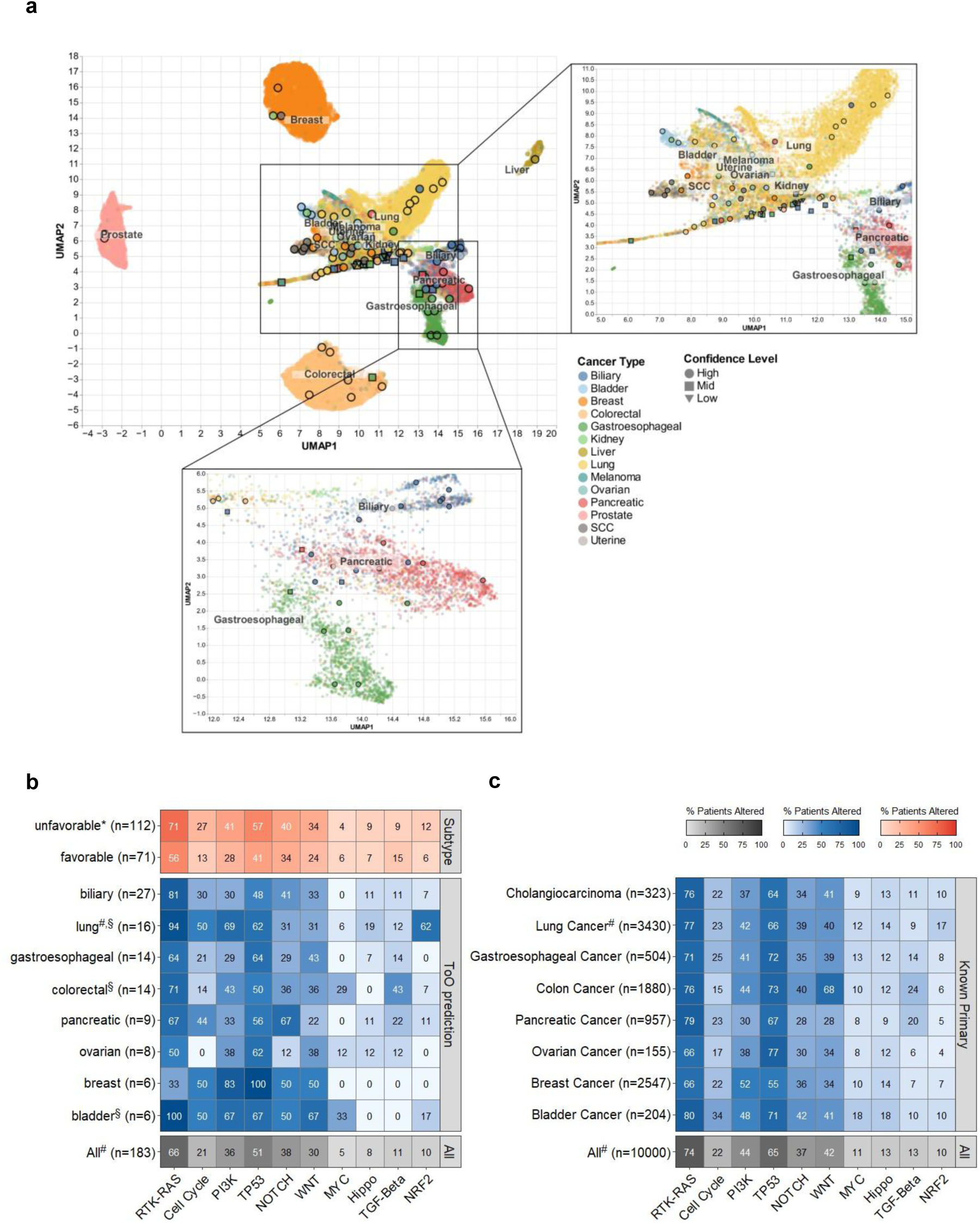
Tissue of origin predictions reflect methylation and mutation features of matching cancer types. **(a)** UMAP of methylation signatures detected in liquid biopsies of CUP patients and patients with a known primary. **(b,c)** Heatmap showing the frequency of patients harboring mutations in one of the ten canonical cancer related pathways across CUP subtypes and ToO predictions, and across cancer types with a known primary, respectively. *: significantly enriched across CUP subtypes using Fisher’s exact test with FDR correction (RTK-Ras: p=0.0395, Cell cycle: p=0.0264, TP53: p=0.0348), §: significantly enriched across ToO predictions using Fisher’s exact test with FDR correction (NRF2: p*=0.0008, MYC: p*=0.0334,), #: significantly enriched between CUP and cancer of known primary using Fisher’s exact test with FDR correction (NRF2 in lung ToO vs lung cancer: p*= 0.0064, TP53 in all CUP vs all cancer of known primary: p*= 0.0064, WNT in all CUP vs all cancer of known primary: p*= 0.0434).

### Lung ToO is associated with NRF2 pathway alterations

Prior studies have shown that alterations in specific cancer-related pathways are significantly associated with specific tumor types [27]. Therefore, we applied the ten canonical pathways profiled by The Cancer Genome Atlas (TCGA) [27] to our CUP cohort. The most frequently altered pathways were RTK-Ras (66%), TP53 (51%), and PI3K (36%), consistent with tissue-based findings in CUP [18]. Additionally, NOTCH (38%) and WNT (30%) signaling pathways were commonly altered in our cohort (Figure 4b). Among patients with unfavorable CUP, cell cycle, TP53, and RTK-RAS pathways were significantly more frequently mutated compared to patients with favorable subtypes, although the differences were not significant after FDR correction. Relative to ToO predictions, we observed significant FDR-adjusted differences in the frequency of NRF2 and MYC pathway alterations (p*=0.0008 and p*=0.0334, respectively; Figure 4b). Alterations in the NRF2 pathway were specifically frequent in patients with a lung ToO prediction (62%), while MYC pathway alterations were frequently observed in patients with a bladder or colorectal ToO prediction. We also compared pathway alteration results from the CUP cohort to cancer data generated with the same assay from patients (n=10,000) with a known primary tumor diagnosis (Figure 4c). Overall, the pattern of pathway alterations was similar between CUP cases and samples with known primary tumor diagnosis. Notably, in our CUP subgroup with a lung ToO prediction, the NRF2 pathway was significantly enriched compared to the lung cancer cohort (p* = 0.0064, Figure 4b,c). As CUP is a metastatic disease, this is in line with NRF2’s role in metastatic lung cancer [28,29]. In addition to the ten canonical pathways, we analyzed DNA damage repair pathways as annotated in the KEGG database (Figure S3c,d). No significant differences were detected by subtypes, ToO predictions, or between CUP and cancer of known primary (Figure S3c).

### Complementary value of combining mutation and methylation profiling

While mutation and methylation profiling have each been established as stand-alone tools for tumor-agnostic and site-specific treatment recommendations in CUP, respectively, they have only been applied independently. To leverage their clinically meaningful information, we combined both modalities in the present study. We applied cancer type-specific therapy implications (OncoKB™) to MTT-predicted cancer types. Actionable gene mutations were detected in 30% of patients with a biliary cancer prediction, 25% with a lung cancer prediction, 14% with a gastroesophageal cancer prediction, and 7% with a colorectal cancer prediction. After incorporating additional tumor-agnostic biomarkers, including TMB-High and MSI-High, 54%, 81%, 21%, and 36% of patients, respectively, were potentially eligible for targeted treatment using cancer type-specific therapies (Figure 5a). Furthermore, combining clinical information with both methylation- and mutation-based ToO predictions and target alterations, we identified potentially therapeutic-relevant clinical information in 137 of 190 (72%) patients (Figure S3e). Importantly, taking all results together, there are two factors that primarily influence the informative value of ctDNA CGP: CUP subtype and status at the time of blood draw. In patients who received ctDNA CGP before treatment, the proportion of patients for whom clinically meaningful information became available increased to 88% (65/74). Furthermore, in 90% of patients with an unfavorable subtype, where the majority of patients still rely on chemotherapy, we could identify potentially therapeutically relevant information (Figure 5b). Therefore, our results show the potential of liquid biopsy approaches for identifying ToO and informing therapeutic strategies for CUP patients, thereby potentially improving the outcome in this prognostically grave disease.

**Figure 5:**
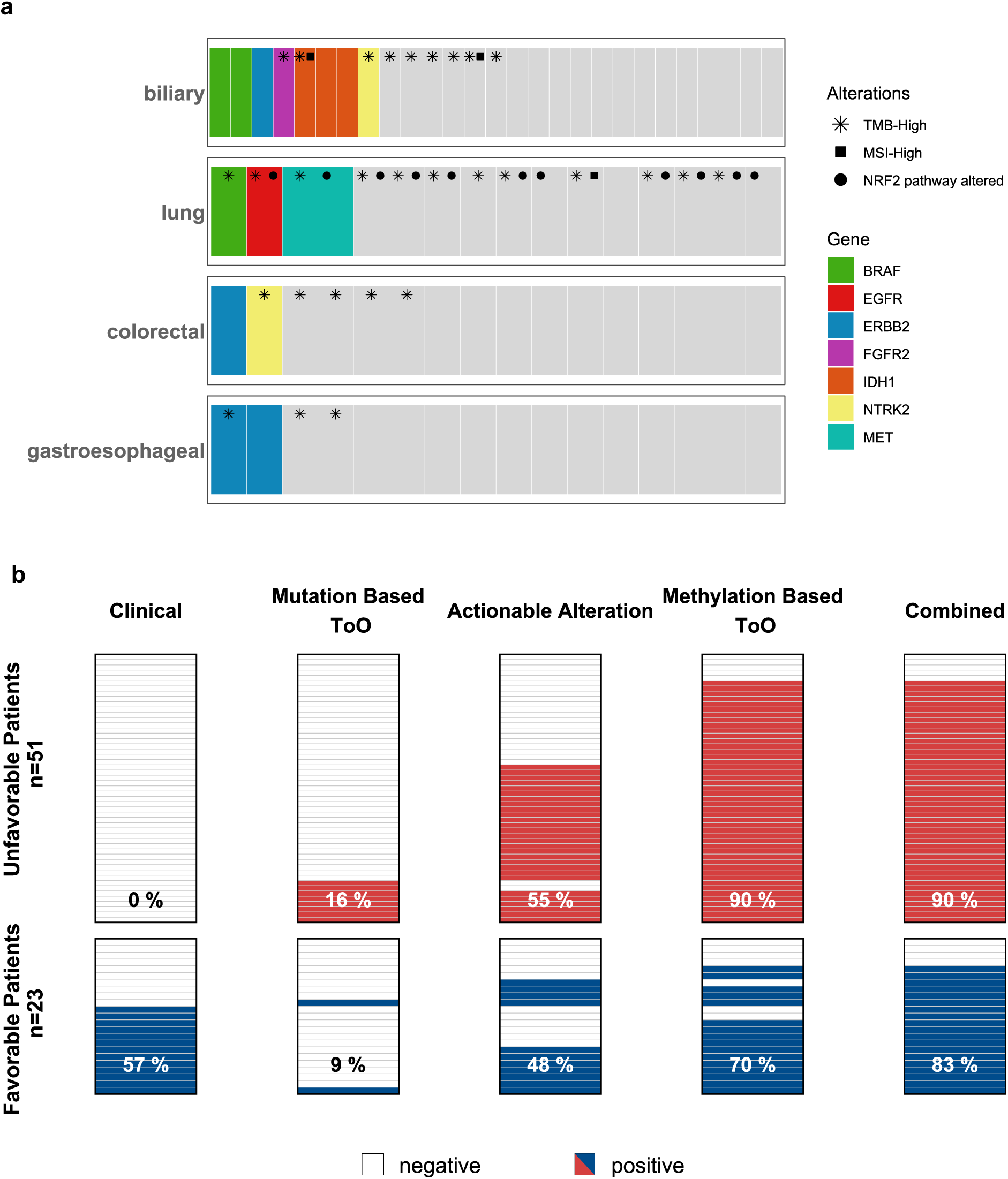
Combining mutation and methylation signatures provides complementary value. **(a)** Tiles plot showing among patients (x-axis) with a positive ToO prediction, tumor type-specific molecular alterations with potential implications for molecularly guided therapy. **(b)** Tiles plot showing the contributions of clinical diagnostic procedure, mutation-based ToO prediction, actionable alteration detection, and methylation-based ToO prediction to clinical therapeutic decision-making in patients analyzed at first diagnosis (n=74).

### ctDNA-based Risk Stratification

Patients were stratified according to EpiTF levels into three groups: undetectable EpiTF, low EpiTF (0–1.5%), and high EpiTF (>1.5%). This classification revealed marked differences in clinical outcome: Patients with high EpiTF had significantly shorter progression-free survival (PFS) and overall survival (OS) compared with those with low or undetectable EpiTF (Figure 6a,b, Log-rank test, p < 0.0001 for both). Median PFS and OS were not reached in the undetectable EpiTF group, with 9.5 and 16.9 months in the low EpiTF group, and only 3.7 and 6.9 months in the high EpiTF group, respectively. Notably, the median PFS observed in the high EpiTF group corresponded to disease progression at first radiographic restaging, indicating a subgroup of patients with particularly aggressive disease. The prognostic value of EpiTF was maintained in subgroup analyses of both unfavorable and favorable CUP patients (Figure 6c–f; unfavorable CUP: PFS, p = 0.0037; OS, p = 0.0064; favorable CUP: PFS and OS, p < 0.0001 for both) as well as in patients analyzed at first diagnosis prior to treatment initiation (Figure S4a,b; PFS, p = 0.0097; OS, p = 0.025). Thus, EpiTF-based stratification consistently identified patient groups with distinct outcomes across different clinical CUP subsets. Comparable results were obtained when patients were stratified into equally sized tertiles according to the number of mutations detected or cfDNA content (number of mutations: Figure S5a-h; cfDNA content: Figure S6a-h).

**Figure 6:**
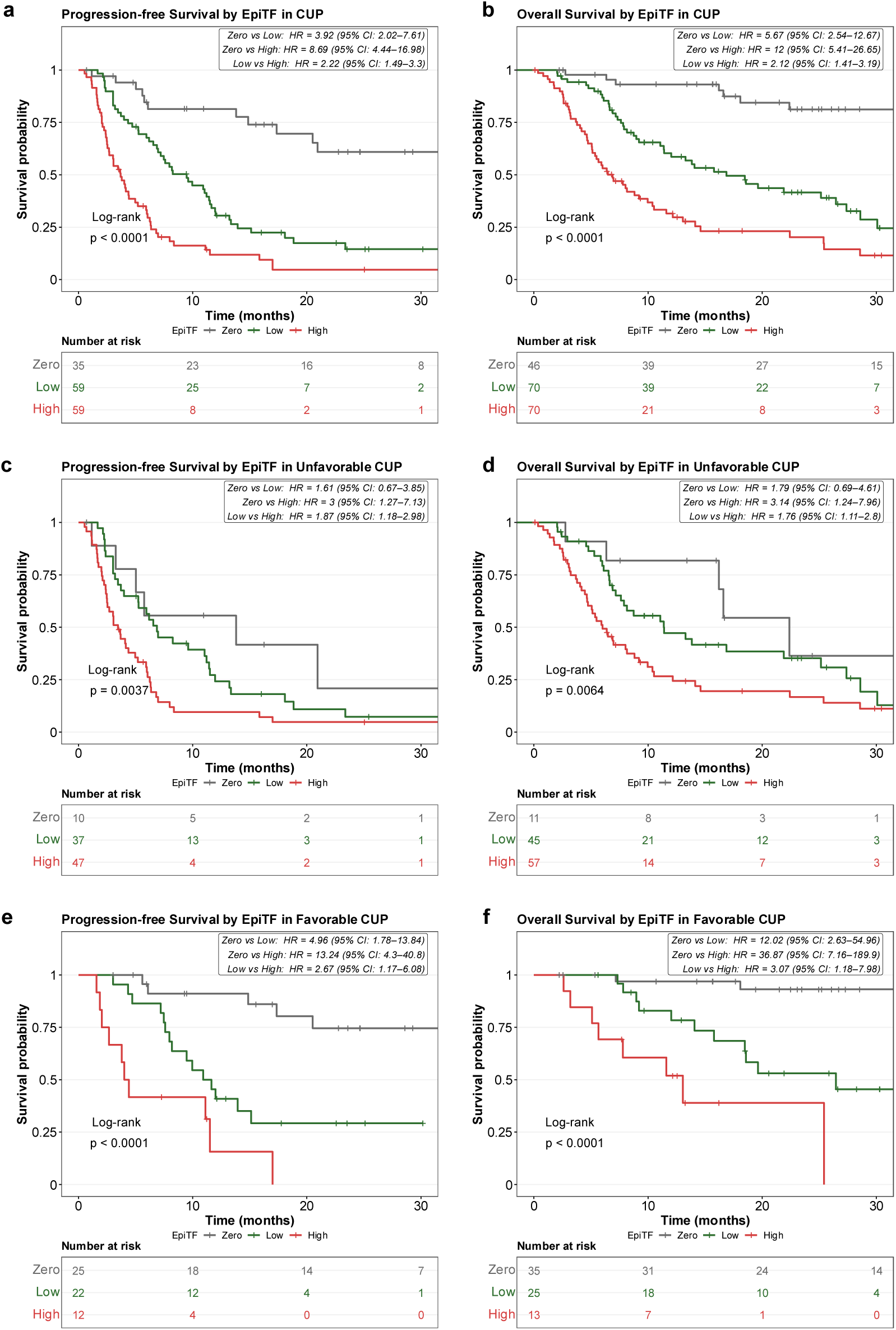
Inferior progression-free survival and overall survival are associated with high tumor fraction. **(a)** Kaplan-Meier plot and risk table of Progression-free survival (PFS) by EpiTF of whole cohort (n=153). Log-Rank test, p<0.0001, HR (95% CI): Zero vs Low: 3.92 (2.02-7.61), Zero vs High: 8.69 (4.44 – 16.98), Low vs High: 2.22 (1.49 - 3.3) **(b)** Kaplan-Meier plot and risk table of overall survival (OS) by EpiTF of whole cohort (n=186). Log-Rank test, p<0.0001, HR (95% CI): Zero vs Low: 5.67 (2.54 - 12.67), Zero vs High: 12 (5.41 – 26.65), Low vs High: 2.12 (1.41 - 3.19). **(c)** Kaplan-Meier plot and risk table of PFS by EpiTF in unfavorable CUP patients (n=94). Log-Rank test, p=0.0037, HR (95% CI): Zero vs Low: 1.61 (0.67-3.85), Zero vs High: 3 (1.27 – 7.13), Low vs High: 1.87 (1.18 - 2.98) **(d)** Kaplan-Meier plot and risk table of OS by EpiTF of unfavorable CUP patients (n=113). Log-Rank test, p=0.0064, HR (95% CI): Zero vs Low: 1.79 (0.69 - 4.61), Zero vs High: 3.14 (1.24 – 7.96), Low vs High: 1.76 (1.11 - 2.8). **(e)** Kaplan-Meier plot and risk table of PFS by EpiTF of favorable CUP patients (n=59). Log-Rank test, p<0.0001, HR (95% CI): Zero vs Low: 4.96 (1.78 – 13.84), Zero vs High: 13.24 (4.3 – 40.8), Low vs High: 2.67 (1.17 – 6.08). **(f)** Kaplan-Meier plot and risk table of OS by EpiTF of favorable CUP patients (n=73). Log-Rank test, p<0.0001, HR (95% CI): Zero vs Low: 12.02 (2.63 – 54.96), Zero vs High: 36.87 (7.16 – 189.9), Low vs High: 3.07 (1.18 – 7.98).

## Discussion

Comprehensive molecular profiling has become increasingly relevant in CUP, a disease characterized by pronounced clinical and biological heterogeneity and limited efficacy of empiric chemotherapy, particularly in patients with unfavorable CUP. In this study, we performed a blood-based analysis of genetic and epigenetic ctDNA features in a large, clinically well-annotated CUP cohort. Using a combined mutation- and methylation-based profiling approach, somatic alterations were detected in 98% (183/187) of patients, actionable biomarkers were identified in 41% (76/187), and methylation-based tissue-of-origin (ToO) prediction was achieved in 65% (122/187). Integrating genomic, epigenomic, and clinical information ultimately provided potentially clinically relevant findings in 72% (137/187) of patients. Importantly, the clinical output was highest in patients with unfavorable CUP and when liquid biopsy was performed at first diagnosis, supporting the potential value of early ctDNA-based comprehensive genomic profiling as an addition to standard tissue-based diagnostics.

The role of ToO-directed therapy in CUP remains controversial. While earlier randomized trials failed to demonstrate a clear survival benefit of site-specific treatment guided by ToO prediction [30,31], the recent FUDAN CUP-001 trial reported improved progression-free survival using a treatment strategy that combined ToO prediction with molecularly guided therapies, including targeted therapy and immunotherapy [7]. As a result, the specific contribution of ToO prediction remains difficult to determine. In contrast, the CUPISCO trial has provided stronger evidence supporting molecularly guided treatment in unfavorable CUP [5]. Taking together, current evidence suggests that ToO information could be more valuable when integrated with actionable genomic findings to inform treatment decisions.

Our findings support an integrated molecular profiling approach. Methylation-based ToO prediction achieved a higher prediction rate than mutation-based prediction using panel sequencing and showed high concordance with clinically defined favorable CUP subsets as well as methylation patterns from cancers of known primary origin. While mutation-based ToO prediction using WES or WGS can be informative when sufficient high-quality tissue is available [32,33], its application in ctDNA is currently limited by costs and sequencing depth requirements. Although ToO prediction alone is unlikely to guide treatment decisions, it can provide valuable context for interpreting actionable alterations. This is particularly relevant because many targeted therapies and immunotherapies remain approved or reimbursed in cancer type-specific settings, and response to the same molecular alteration may differ depending on tumor origin [34]. Combining ToO information with genomic profiling may therefore improve treatment stratification and expand therapeutic options in selected CUP patients.

A few liquid biopsy-based studies on mainly a limited number of CUP have been published before, either determining ToO or detecting actionable alterations [10,15,35–37]. Our study, together with previous liquid biopsy-based studies, highlights the complementary value of liquid biopsy compared with tissue-based profiling. Tissue availability is frequently limited in CUP due to extensive diagnostic work-up, small biopsies, or the need for repeated immunohistochemical and molecular analyses [4]. Liquid biopsy may overcome some of these limitations by enabling minimally invasive diagnostics, capturing genomic heterogeneity across multiple metastatic sites, and allowing repeated sampling during the disease course [16,17]. In our cohort, ctDNA analysis provided actionable information beyond tissue sequencing in a subset of patients, particularly when performed early, supporting its potential role in the initial diagnostic and therapeutic assessment of CUP.

From a biological perspective, mutational analysis of cancer-related pathways showed a notably high frequency of NRF2 pathway alteration in patients with a lung ToO, consistent with the frequent NRF2 pathway alterations observed in lung cancer [27]. This is particularly relevant in a metastatic disease setting such as CUP, as NRF2 activation is associated with metastasis in lung cancer [28]. Importantly, drugs targeting this pathways have been tested in squamous non-small cell lung cancer with an overall response rate of 25% [38].

In addition to its diagnostic and therapeutic implications, ctDNA profiling provided clinically relevant prognostic information. EpiTF was strongly associated with progression-free and overall survival across the overall cohort and within clinically relevant subgroups, including both favorable and unfavorable CUP as well as patients analyzed at first diagnosis. The particularly poor outcome of patients with high EpiTF, whose median PFS approximately corresponded to progression at first radiographic restaging, identifies a subgroup with highly aggressive disease biology and a high risk for early treatment failure. This finding suggests that ctDNA quantification could help identify patients who require closer monitoring, early treatment adaptation, or inclusion in interventional trials.

Although this represents one of the largest liquid biopsy cohorts in CUP analyzed to date, the cohort reflects the substantial clinical and biological heterogeneity of this entity. This limited the statistical power for detailed analyses of individual favorable CUP subsets. In addition, although methylation-based ToO prediction achieved a high overall prediction rate, some categories, particularly squamous cell carcinoma, remain insufficiently specific to reliably predict a distinct anatomical primary site and may therefore provide limited guidance for site-specific treatment selection. Finally, because of the observational study design, the impact of ctDNA-guided therapeutic strategies on clinical outcome cannot be directly assessed and will require prospective validation in interventional trials.

In conclusion, this study demonstrates that integrated mutation and methylation profiling from plasma samples provides clinically relevant diagnostic, prognostic, and potentially therapeutic information in CUP. The combined assessment of actionable alterations, methylation-based ToO prediction, and ctDNA-derived tumor fraction may be particularly valuable in unfavorable CUP and when applied early in the disease course. These findings support the incorporation of liquid biopsy into future prospective studies evaluating molecularly guided management strategies for CUP.

## Material and Methods

### Patients

Blood from 264 patients with suspected CUP who presented to Heidelberg University Hospital between July 2021 and December 2024 was collected (Figure 1a). Patients with a confirmed CUP diagnosis per the European Society of Medical Oncology (ESMO) clinical practice guidelines were eligible for inclusion in this study [4]. In cases where relapsed antecedent malignancies could be misidentified as CUP, comparative tissue sequencing was performed to confirm the CUP diagnosis, as previously described [39]. Further exclusions of suspected CUP cases were most often due to incomplete diagnostic processes at the start of the study. The eligible cohort of 190 CUP patients included those with newly diagnosed CUP and those who presented during treatment, at disease progression, or at follow-ups. The distinction between the unfavorable subtype and favorable subsets followed current ESMO guidelines and was based on histology, immunohistochemistry, pattern and magnitude of metastatic spread, and additional diagnostic work-up, informed by medical history and symptoms. The median age of the cohort consisting of 110 (58%) female and 80 (42%) male patients was 63 years (range: 23-85). The majority of patients, 115 (61%), were diagnosed with unfavorable CUP. The favorable CUP subtype (n=75, 39%) was more common than in the general population [4], likely due to overrepresentation within a tertiary cancer center. Histopathological examination of tissue biopsies revealed mainly adenocarcinomas (n=129, 68%) and squamous cell carcinomas (n=39, 21%). Seventy-four (39%) samples were collected at first diagnosis, 59 (31%) during treatment, 31 (16%) during disease progression, and 26 (14%) at follow-up. The cohort displayed a wide range of affected organs, with lymph nodes being the most prevalent (n=98, 52%). Detailed patient cohort information is given in Supplementary Table S2. For 127 patients, panel- or whole-genome/whole-transcriptome-sequencing (TSO500: n=58, WES/WTS: n=69) of the tumor tissue was performed at the pathology department of the University Hospital Heidelberg to guide individualized treatment. Clinical data were collected from patients’ medical records. All patients provided written informed consent. The study was conducted in accordance with Good Clinical Practice guidelines and the Declaration of Helsinki and was approved by the Ethics Committee of the University of Heidelberg (S-662/2021).

### Sample collection

Whole blood was collected in Cell-Free DNA BCT tubes (Streck, La Vista, NE, USA) and plasma isolation occurred within 96 h using sequential centrifugation at 1800 g and 4700 g. Subsequently the plasma was stored at -80°C until further analysis. Samples were batch shipped on dry ice to Guardant Health (Palo Alto, CA, USA) for the isolation of cell free DNA (cfDNA) and next generation sequencing (NGS) using Guardant360 Liquid.

### Isolation of cfDNA and Sequencing

cfDNA was extracted from plasma samples as previously described [40]. Extracted cfDNA was analyzed using the Guardant360 Liquid assay, which simultaneously profiles genomic alterations and DNA methylation from plasma-derived cfDNA at single-molecule resolution [41]. The assay interrogates >700 genomic biomarkers and >20,000 epigenomic regions. Genomic alterations included in subsequent analyses comprised single nucleotide variants (SNVs), insertions and deletions (indels), copy number alterations (including amplifications and losses), gene fusions, and microsatellite instability (MSI).

Tissue-of-origin prediction was performed using Molecular Tumor Typing (MTT), a methylation-based tissue-of-origin classifier integrated into the Guardant360 Liquid assay. MTT comprises an ensemble of 14 regression models (one for each cancer type; Supplementary Table S3) based on >3,000 differentially methylated regions (DMRs). For each sample, a model score was calculated by each regression model. Based on the predicted sex, sex-incompatible cancer types were excluded, including prostate cancer for females and breast, ovarian and uterine cancers for males. The cancer types with the highest and second highest scores were reported as primary and secondary predictions, respectively.

### Circulating tumor cell (CTC) enrichment and enumeration

For 158 patients, simultaneously with the plasma sample, 7.5 mL whole blood was collected in CellSave preservative tubes (Menarini Silicon Biosystems, Florence, Italy). CTCs were isolated from blood samples using CellSearch® technology (Menarini Silicon Biosystems, Florence, Italy) as previously described [23,42]. In brief, blood samples were stored at room temperature and were analyzed within 96 hours of collection using the CellSearch assay according to the manufacturer’s instructions. Cells are enriched using antibodies to epithelial cell adhesion molecule (EpCAM) attached to a ferrofluid. Subsequently, cells of epithelial origin are immunomagnetically separated from cellular blood components. Afterwards, the enriched cells were stained with fluorescently labelled antibodies against cytokeratins including CK8, CK18 and CK19, as well as CD45 and DAPI to differentiate between debris, hematopoietic cells and CTCs. CTCs were defined as nucleated cells expressing cytokeratins but not CD45. The images generated by the CellSpotterTM Analyzer, a semiautomated fluorescence microscope, were evaluated by experienced operators.

### Downstream analysis and graphical representation

The downstream analyses described below was performed with R (version 4.5.2) [43] in RStudio (version 2026.04.0+526) using mainly tidyverse (version 2.0.0) [44]. Genomic analysis were carried out using Bioconductor repository (version 2.70.0) [45] including maftools (version 2.26.0) [46] and VariantAnnotation (version 1.56.0) [47] packages. Graphical representation was accomplished using extensions to ggplot (part of tidyverse), namely patchwork (version 1.3.2), ggnewscale (version 0.5.2), ggpubr (version 0.6.3), and ComplexUpset (version 1.3.3) [48]. Sankey Diagram was created using sankeyD3plus (version 0.2) [49]. Survival analysis was accomplished using R package survival (version 3.8-6) [50], Kaplan Meier curves were created using survminer (version 0.5.1) [51].

### Identification of actionable alterations

Actionable alterations were retrieved from the FDA-recognized MSK’s Precision Oncology Knowledge Base OncoKB™ [24,25]. Actionable alterations within levels of evidence 1 and 2 were reported. Calling of actionable alterations was validated by an experienced physician. To compare mutually detectable actionable alterations between liquid and tissue biopsy, mutations detected in one biopsy specimen were filtered to include only those that were detectable by the corresponding alternative assay.

### Pathway analyses

The ten canonical pathways described by Sanchez-Vega et al., 2018 [27] were exported from maftools package. Genes that belong to DNA repair pathways were retrieved from Kyoto Encyclopedia of Genes and Genomes (KEGG) using pathway identifiers hsa03410, hsa03420, hsa03430, hsa03440, hsa03450, hsa03460 in R package KEGGREST (version 1.50.0) [52]. Pathway information is described in Supplementary Table S4. For each pathway the proportion of patients that harbor at least one of the following somatic alterations, non-synonymous SNV, Indel, high-level amplification, homologous deletion, or Fusion in at least one of the described genes was determined. Mutation data from patients with cholangiocarcinoma, lung cancer, gastroesophageal cancer, colon cancer, pancreatic cancer, ovarian cancer, breast cancer, or bladder cancer that underwent Guardant360 Liquid analysis were received from the InfinityAI Data Library (Guardant Health, Palo Alto, CA). Pathway analysis was accomplished as described above.

### Unsupervised clustering of CUP and cancer with known primary

A reference sample of 34,911 patients with advanced stage cancer was compiled from InfinityAI Data Library. The Uniform Manifold Approximation and Projection (UMAP) projection analysis uses scikit-learn (v1.0.0)’s Principal Component Analysis (PCA) and the umap-learn (v0.5.3) library to reduce the high-dimensional differential methylated region (DMR) data to a 2D embedding for visualization. A PCA model is fit on the reference samples retaining principal components (PCs) 2–16 and PC1 is excluded due to its high correlation with tumor fraction. Test samples (CUP cohort) are projected through the same frozen PCA. A UMAP model (15 neighbors, min distance 0.3, Euclidean metric) is then fit on the reference PCA coordinates, and test samples are projected into that fixed embedding. Altair (v4.2.0) package is used to visualize the UMAP embeddings.

### Survival Analyses

Survival analyses were conducted for epigenetic tumor fraction (EpiTF), cell-free DNA (cfDNA), and number of somatic mutations. Based on the EpiTF thresholds of > 0 and < 1.5%, patients were stratified into EpiTF-negative, EpiTF-low, and EpiTF-high. The median of all EpiTF-positive patients was chosen as the cut-off. For PFS 35, 59, and 59 patients were analyzed per strata; for OS, 46, 70, and 70 patients. The thresholds of 0 and 1.5% were inherited for all subgroup analyses. Patients were categorized based on the cfDNA into three equally sized tertiles: low (≤ 17.74 ng), medium (> 17.74 & ≤ 30 ng) and high (> 30 ng). Furthermore, patients were categorized based on the number of non-synonymous somatic mutations into three tertiles: low (0-6 mutations), medium (7-18 mutations), and high (19 or more mutations). Progression-free survival (PFS) was defined as the time from the initiation of baseline therapy, the therapy closest to the first liquid biopsy, until progression, death, relapse, or censoring at the end of follow-up. OS was measured from the date of the first ctDNA collection until death or the end of follow-up.

### Statistical Analyses

Correlations among continuous data were assessed by Spearman’s rank correlation coefficient. Fisher’s exact test was performed to test differences between groups for nominally scaled and dichotomous data. Cramér’s V was calculated following Fisher’s exact test to assess the effect size for large sample sizes. Group comparisons of continuously scaled data were carried out using Kruskal-Wallis, following post-hoc Dunn’s test for multiple comparisons or the Wilcoxon rank sum test. The Kaplan–Meier method was used to estimate median progression-free survival and median OS for each group. Progression-free survival and OS were compared between groups by the stratified Log-Rank test, with hazard ratios (HRs) and corresponding 95% confidence intervals (CIs) estimated by use of a stratified Cox proportional hazards model. False discovery rate (FDR) correction was applied for multiple comparisons, and adjusted p values are reported as p*. p- and p*-values <0.05 were considered statistically significant.

## Declarations

### Ethics approval and consent to participate

The study was approved by the Ethics Committee of the University of Heidelberg (S-662/2021). All patients provided written informed consent.

