## Supplementary Data for "Comprehensive ctDNA Profiling Enables Tissue-of-Origin Prediction and Actionable Biomarker Detection in Cancer of Unknown Primary"

### Supplemental Data

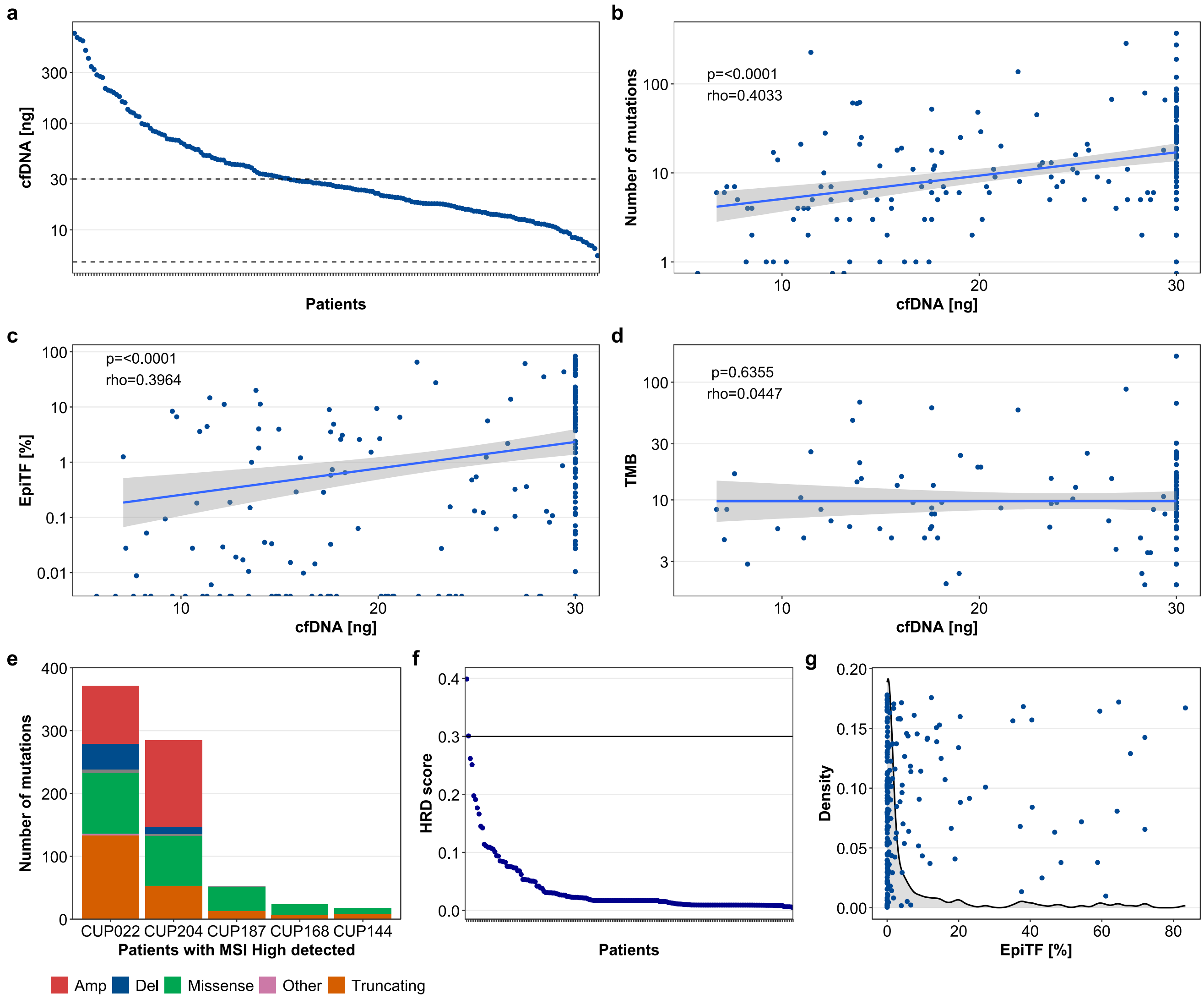


**Figure S1: Highly sensitive ctDNA panel. (a)** Point graph showing the amount of cell free DNA (cfDNA) isolated from plasma, patients ordered by decreasing cfDNA yield. Dashed lines representing the minimal recommended and the maximal input of 5 and 30 ng, respectively, of cfDNA for the assay. **(b,c,d)** Correlation of the number of mutations, EpiTF or TMB detected to the cfDNA input, Spearman's rank correlation coefficient; number of mutation vs cfDNA: p<0.0001, rho=0.4022; EpiTF vs cfDNA: p<0.0001, rho=0.3964; TMB vs cfDNA: p=0.5591, rho=0.0546. **(e)** Barplot showing the number of mutations of patients with MSI High detected colored by mutation type, Amp amplification, Del deletion. **(f)** Point graph showing the HRD score determined by the HRD module, patients ordered by decreasing cfDNA yield. Line representing the threshold for HRD positivity. **(g)** Density plot showing the distribution of patients by EpiTF using Kernel density estimates (curve) overlaid with individual patient EpiTF scores (x-axis) distributed random along the y-axis (points).

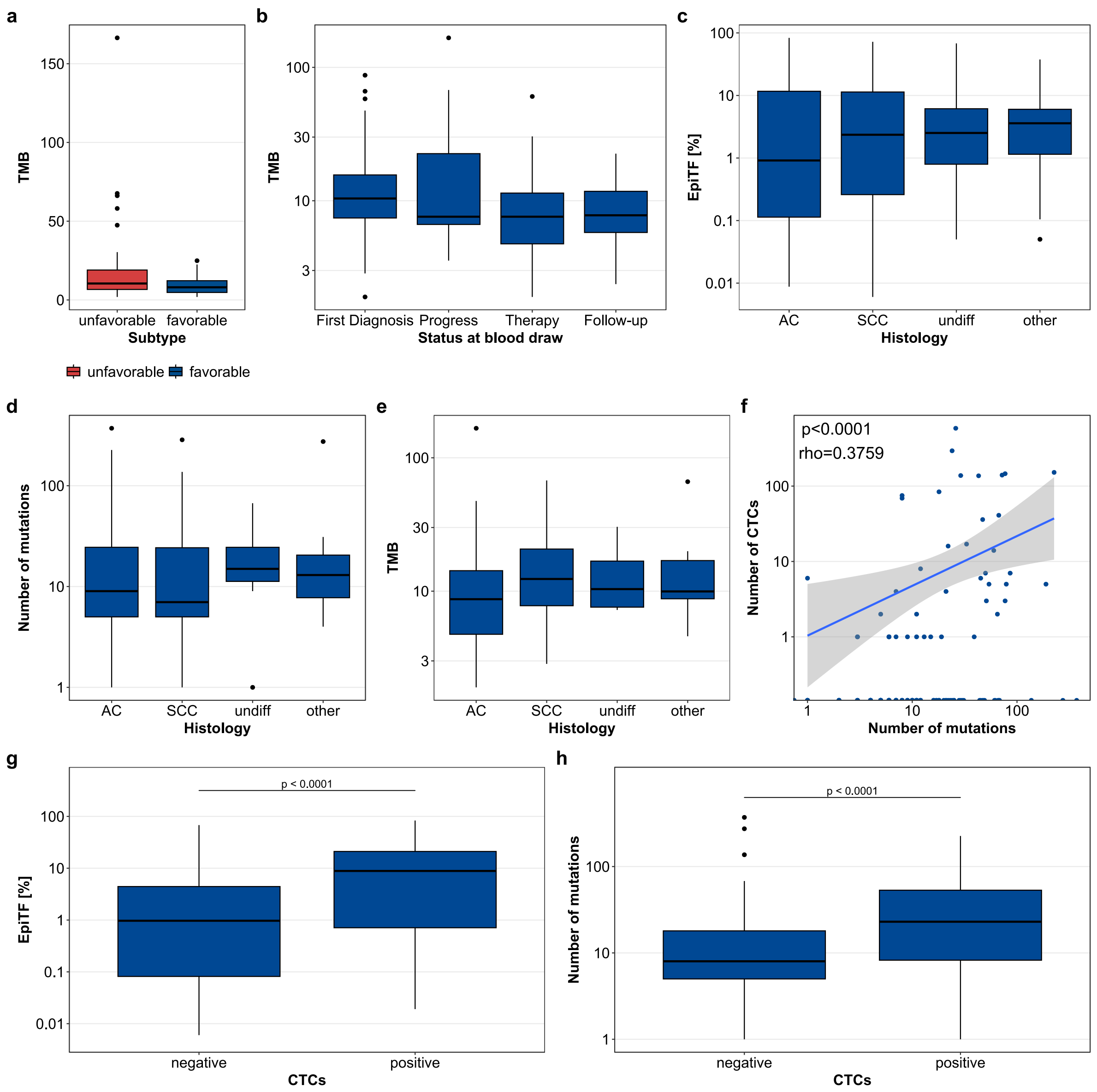


**Figure S2: Correlation of clinical features with ctDNA. (a,b)** Boxplots depicting no correlation between TMB and subtype or status at blood draw, respectively. Subtype: Wilcoxon rank sum test, p=0.0843; status at blood draw: Kruskal-Walli's rank sum test, p=0.2069. **(c,d,e)** Boxplots depicting no association between Histology (AC: adenocarcinoma, SCC: squamous-cell carcinoma, undiff: undifferentiated carcinoma) and EpiTF, number of mutations or TMB, respectively. Kruskal-Walli's rank sum test, EpiTF: p=0.1951,number of mutations: p=0.3195, TMB: p=0.1732. **(f)** Correlation of the number of CTCs with number of mutations, Spearman's rank correlation coefficient, p<0.0001, rho=0.3776. **(g,h)** Boxplots showing the association between CTC status and EpiTF or number of mutations, respectively. Wilcoxon's rank sum test, EpiTF: p<0.0001, number of mutations: p<0.0001. For all boxplots, the center line indicates the median, boxes represent the 1st and 3rd quartiles, whiskers extend to 1.5 x the interquartile range (IQR), and points depict outliers of 1.5xIQR.

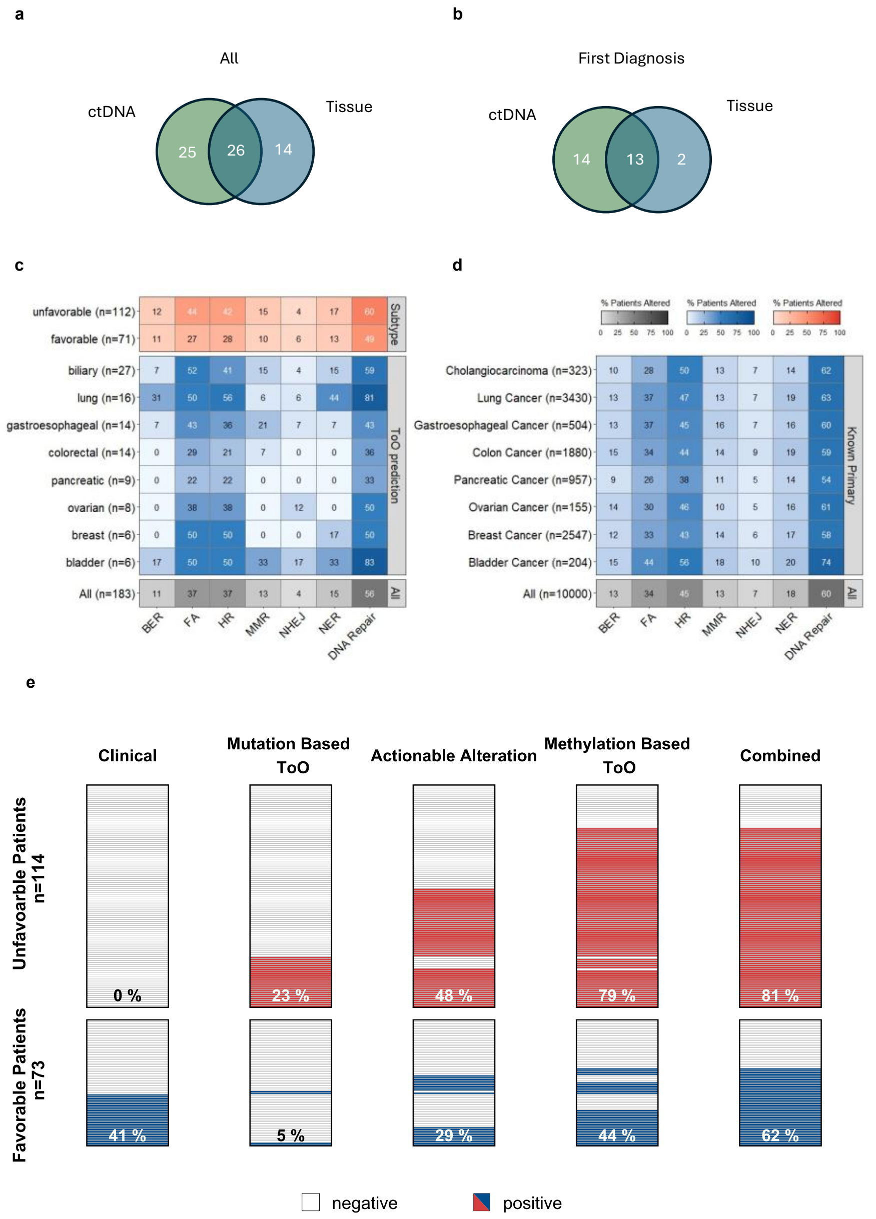


**Figure S3: Comparison of patients with actionable alterations between ctDNA and tissue, DNA repair pathways and clinical implications. (a, b)** Venn-diagrams showing the number of patients harboring mutual actionable alterations detected in liquid and tissue biopsy for all patients (n=65) and patients that received both sequencing approaches before treatment (first diagnosis n=29), respectively. **(c,d)** Heatmap showing the frequency of patients harboring mutations in one of the KEGG DNA repair pathways across subtype and ToO predictions and across cancer types with a known primary, respectively. **(e)** Tilesplot showing the different contributions to clinical therapeutic decision making for patients (y-axis) of clinical diagnostic procedure, mutation-based ToO prediciton, actionable alteration detection, methylation-based ToO prediction, for all patients (n=183).

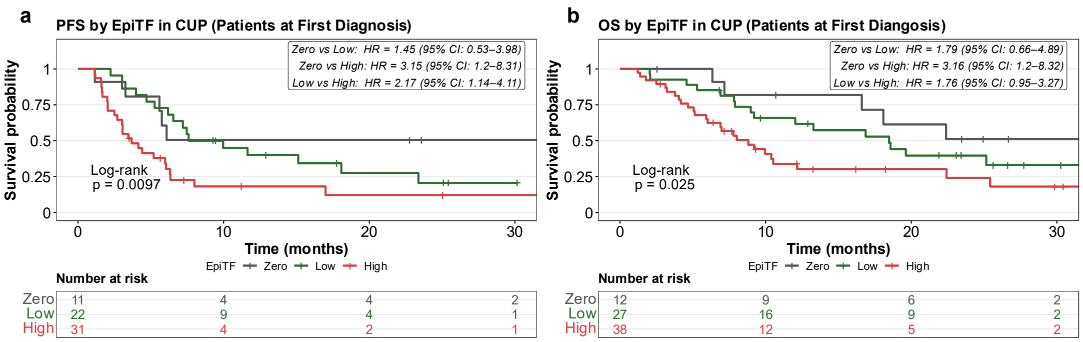


**Figure S4: (a)** Kaplan-Meier plot and risk table of progression-free survival (PFS) by EpiTF of patients at first diagnosis prior to treatment initiation (n=64). Log-Rank test, p=0.0097. **(b)** Kaplan-Meier plot and risk table of overall survival (OS) by EpiTF of patients at first diagnosis prior to treatment initiation (n=77). Log-Rank test, p=0.025.

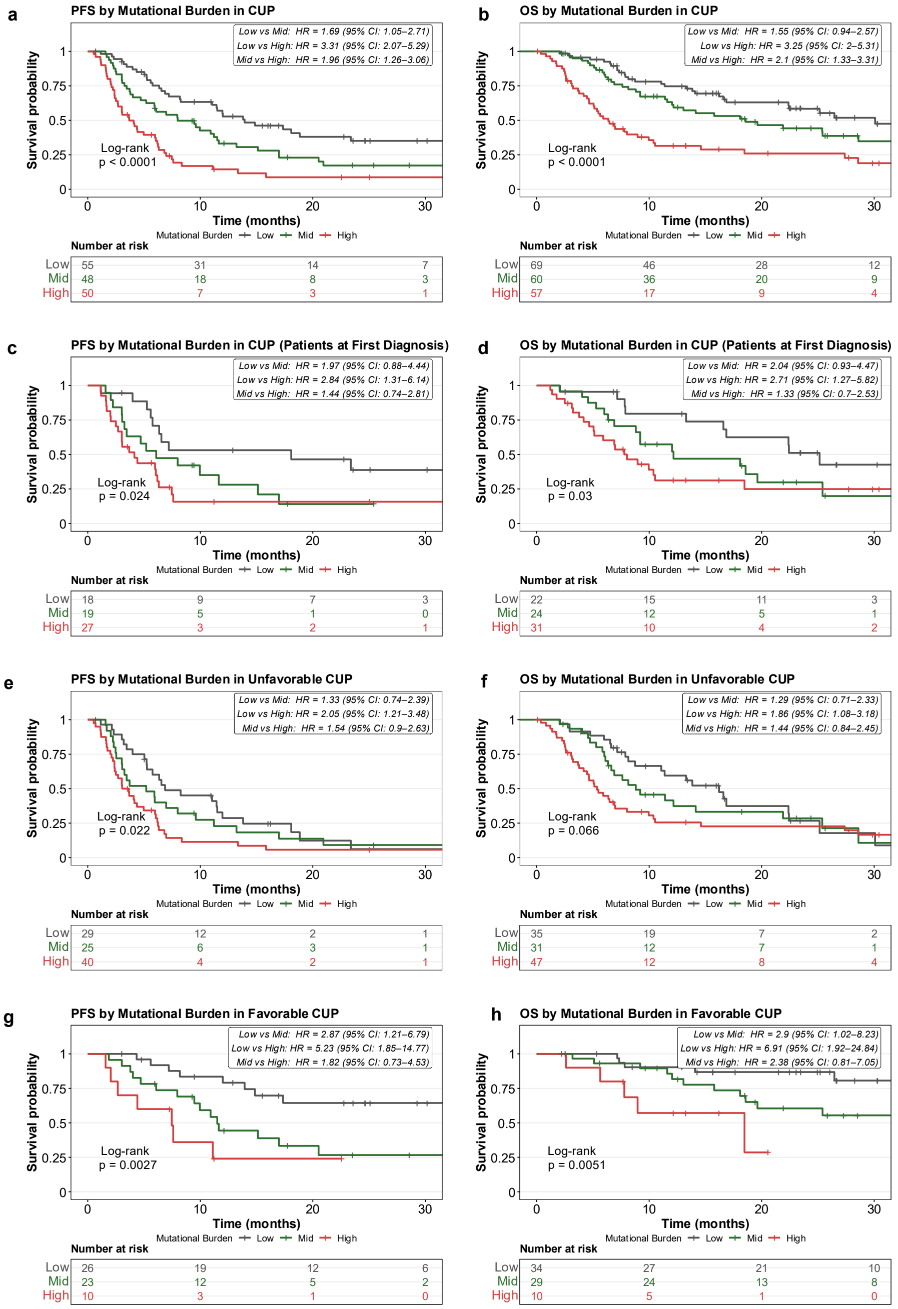


**Figure S5: (a,b)** Kaplan-Meier plot and risk table of progression-free survival (PFS) and overall survival (OS) by mutational burden (number of mutations) of whole cohort (PFS: n=153, OS: n=186), respectively. Log-Rank test, p<0.0001 for both **(c,d)** Kaplan-Meier plot and risk table of PFS and OS by mutational burden at first diagnosis prior to treatment initiation (PFS: n=64, OS: n=77), respectively. Log-Rank test, PFS: p=0.024, OS: p=0.003. **(e,f)** Kaplan-Meier plot and risk table of PFS and OS by mutational burden of unfavorable CUP (PFS: n=94, OS: n=113), respectively. Log-Rank test, PFS: p=0.022, OS: p=0.066. **(g,h)** Kaplan-Meier plot and risk table of PFS and OS by mutational burden of favorable CUP (PFS: n=59, OS: n=73), respectively. Log-Rank test, PFS: p=0.0027, OS: p=0.00051.

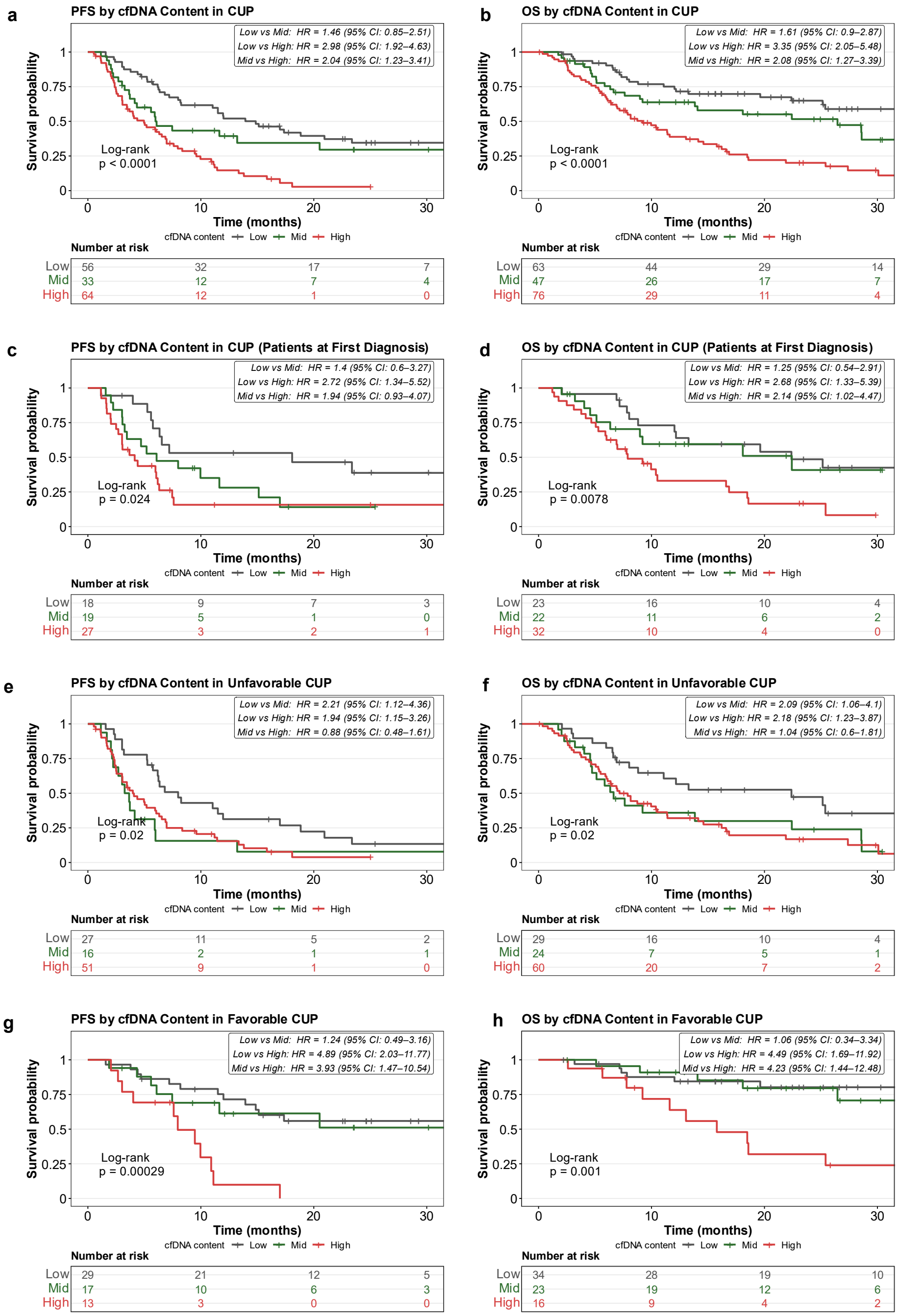


**Figure S6: (a,b)** Kaplan-Meier plot and risk table of progression-free survival (PFS) and overall survival (OS) by cfDNA input of whole cohort (PFS: n=153, OS: n=186), respectively. Log-Rank test, p<0.0001 for both **(c,d)** Kaplan-Meier plot and risk table of PFS and OS by cfDNA input at first diagnosis prior to treatment initiation (PFS: n=64, OS: n=77), respectively. Log-Rank test, PFS: p=0.024, OS: p=0.0078. **(e,f)** Kaplan-Meier plot and risk table of PFS and OS by cfDNA input of unfavorable CUP (PFS: n=94, OS: n=113), respectively. Log-Rank test, p=0.02 for both. **(g,h)** Kaplan-Meier plot and risk table of PFS and OS by cfDNA input of favorable CUP (PFS: n=59, OS: n=73), respectively. Log-Rank test, PFS: p=0.00029, OS: p=0.001.

**Supplementary Table S1:** Concordance of mutation-based and methylation-based ToO prediction

| Patient_ID | OncoNPC prediction (mutation) | confidence_score OncoNPC | Clinical Subtype | MTT prediction (methylation) | confidence_score MTT | Correlation |
|---|---|---|---|---|---|---|
| CUP157 | Non-Small Cell Lung Cancer | 0.945 | unfavorable | | | New ToO |
| CUP256 | Non-Small Cell Lung Cancer | 0.888 | unfavorable | | | New ToO |
| CUP216 | Non-Small Cell Lung Cancer | 0.998 | unfavorable | lung | 0.9993 | Concordant |
| CUP195 | Non-Small Cell Lung Cancer | 0.993 | unfavorable | lung | 0.9999 | Concordant |
| CUP086 | Non-Small Cell Lung Cancer | 0.989 | single-site | lung | 0.9993 | Concordant |
| CUP211 | Pancreatic Adenocarcinoma | 0.986 | unfavorable | pancreatic | 0.9993 | Concordant |
| CUP093 | Non-Small Cell Lung Cancer | 0.984 | unfavorable | lung | 0.9844 | Concordant |
| CUP097 | Non-Small Cell Lung Cancer | 0.976 | unfavorable | lung | 0.9447 | Concordant |
| CUP230 | Colorectal Adenocarcinoma | 0.967 | unfavorable | colorectal | 0.9993 | Concordant |
| CUP116 | Non-Small Cell Lung Cancer | 0.922 | unfavorable | lung | 0.7663 | Concordant |
| CUP118 | Non-Small Cell Lung Cancer | 0.898 | unfavorable | lung | 0.9999 | Concordant |
| CUP075 | Non-Small Cell Lung Cancer | 0.884 | unfavorable | lung | 0.9993 | Concordant |
| CUP123 | Esophagogastric Adenocarcinoma | 0.875 | unfavorable | gastroesophageal | 0.9993 | Concordant |
| CUP233 | Non-Small Cell Lung Cancer | 0.832 | unfavorable | lung | 0.9928 | Concordant |
| CUP257 | Invasive Breast Carcinoma | 0.829 | single-site | breast | 0.9997 | Concordant |
| CUP158 | Non-Small Cell Lung Cancer | 0.814 | unfavorable | lung | 0.9275 | Concordant |
| CUP193 | Invasive Breast Carcinoma | 0.825 | breast-like | breast | 0.9963 | Concordant |
| CUP028 | Non-Small Cell Lung Cancer | 0.971 | unfavorable | scc | 0.9999 | Concordant |
| CUP201 | Non-Small Cell Lung Cancer | 0.966 | unfavorable | scc | 0.9993 | Concordant |
| CUP247 | Colorectal Adenocarcinoma | 0.932 | unfavorable | gastroesophageal | 0.8851 | clinically often not distinguishable |
| CUP168 | Pancreatic Adenocarcinoma | 0.842 | unfavorable | biliary | 0.9995 | clinically often not distinguishable |
| CUP223 | Non-Small Cell Lung Cancer | 0.955 | unfavorable | scc | 0.9993 | Discordant |
| CUP145 | Non-Small Cell Lung Cancer | 0.811 | unfavorable | scc | 0.9991 | Discordant |
| CUP225 | Colorectal Adenocarcinoma | 0.805 | unfavorable | scc | 0.9993 | Discordant |
| CUP220 | Melanoma | 0.947 | unfavorable | gastroesophageal | 0.9993 | Discordant |
| CUP100 | Melanoma | 0.940 | unfavorable | scc | 0.7356 | Discordant |
| CUP163 | Bladder Urothelial Carcinoma | 0.879 | colon-like | colorectal | 0.9999 | Discordant |
| CUP106 | Melanoma | 0.847 | unfavorable | bladder | 0.9980 | Discordant |
| CUP098 | Non-Small Cell Lung Cancer | 0.844 | unfavorable | bladder | 0.9994 | Discordant |
| CUP143 | Non-Small Cell Lung Cancer | 0.821 | unfavorable | biliary | 0.9999 | Discordant |

**Supplementary Table S2:** Patient cohort

| | |
|---|---|
| **Age at blood draw (median, range)** | 63 (23-85) |
| **Gender (female/male)** | 110 (57.9%) / 80 (42.1%) |
| **Subtype (Unfavorable / Favorable)** | 115 (60.5%) / 75 (39.5%) |
| Single-Site/ Oligometastatic | 45 (23.7%) |
| Colon-like | 22 (11.6%) |
| Head-and-Neck-like | 3 (1.6%) |
| Breast-like | 3 (1.6%) |
| Ovary-like | 2 (1.1%) |
| **Histology** | |
| Adenocarcinoma | 129 (67.9%) |
| Squamous cell carcinoma | 39 (20.5%) |
| Undifferentiated carcinoma | 10 (5.3%) |
| Other | 12 (6.3%) |
| **Number of affected organs (0/1/2/3/4+)** | 30 (15.8%) / 68 (35.8%) / 44 (23.2%) / 25 (13.2%) / 24 (12.6%) |
| **Organs affected** | |
| No organs affected | 30 (15.8%) |
| Lymph nodes | 98 (51.6%) |
| Liver | 45 (23.6%) |
| Lung | 45 (23.6%) |
| Peritoneum or pleura | 57 (30%) |
| Soft tissue | 22 (11.6%) |
| Other | 30 (15.8%) |
| **Time Point of Baseline Sample (at first diagnosis / at disease progression / under therapy / at follow-up)** | 74 (38.9%) / 33 (17.4%) / 58 (30.5%) / 25 (13.2%) |

**Supplementary Table S3:** Cancer types for which a regression model is integrated in the Molecular Tumor Typing classifiert.

| **Cancer types** |
|---|
| Bladder |
| Breast |
| Biliary (cholangiocarcinoma) |
| Colorectal |
| Endometrial/uterine |
| Gastroesophageal |
| Liver |
| SCC (Squamous Cell Carcinoma) |
| Lung |
| Melanoma |
| Ovarian |
| Pancreatic |
| Prostate |
| Kidney (RCC) |
